# Activated PI3Kδ specifically perturbs mouse Treg homeostasis and function leading to immune dysregulation

**DOI:** 10.1101/2023.12.21.569665

**Authors:** Akhilesh K. Singh, Fahd Al Qureshah, Travis Drow, Baidong Hou, David J Rawlings

## Abstract

Foxp3^+^ regulatory T cells (Treg) are required for maintaining immune tolerance and preventing systemic autoimmunity. PI3Kδ is required for normal Treg development and function. However, the impacts of dysregulated PI3Kδ signaling on Treg function remain incompletely understood. Here, we used a conditional mouse model of activated PI3Kδ syndrome (APDS) to investigate the role of altered PI3Kδ signaling specifically within the Treg compartment. Aged mice expressing a PIK3CD gain-of-function mutation (aPIK3CD) specifically within the Treg compartment exhibited weight loss and evidence for chronic inflammation as demonstrated by increased memory/effector CD4^+^ and CD8^+^ T cells with enhanced IFN-γ secretion, spontaneous germinal center responses and production of broad-spectrum autoantibodies. Intriguingly, aPIK3CD facilitated Treg precursor development within the thymus and an increase in peripheral Treg numbers. Peripheral Treg, however, exhibited an altered phenotype including increased PD1 expression and reduced competitive fitness. Consistent with these findings, Treg specific-aPIK3CD mice mounted an elevated humoral response following immunization with a T-cell dependent antigen, that correlated with a decrease in follicular Treg. Taken together, these findings demonstrate that an optimal threshold of PI3Kδ activity is critical for Treg homeostasis and function, suggesting that PI3Kδ signaling in Treg might be therapeutically targeted to either augment or inhibit immune responses.

## Introduction

Regulatory T cells (Treg) defined by expression of the transcription factor Foxp3 are a specialized subset of T lymphocytes that are required for normal immune homeostasis (Hori et al., 2003). Treg play a central role in maintaining immune tolerance and preventing autoimmune responses, and Treg deficiency or perturbation can lead to autoimmunity and systemic polyclonal lymphoproliferation (Fontenot et al., 2003). The generation, function, and stability of Treg are regulated by a complex interplay of signaling pathways (Huynh et al., 2014). Thus, understanding signals that govern Treg differentiation and function is crucial for defining immune tolerance mechanisms and represent potential therapeutic strategies to modulate Treg responses and restore immune balance in autoimmune and inflammatory disorders.

The phosphoinositide 3-kinase (PI3K) pathway plays a crucial role in biological processes in nearly all cell types. There are three classes of PI3K in mammals and Class I PI3K molecules play the most critical roles in immune cells. Class I PI3Ks are heterodimeric proteins composed of a catalytic p110 subunit and a regulatory subunit. Among Class I PI3Ks, p110δ expression is restricted to immune cells and has distinct functions in adaptive immunity. In T cells, PI3Kδ can be activated by multiple receptors and co-receptors, including the T-cell receptor (TCR), CD28, inducible T-cell co-stimulator (ICOS), and cytokine and chemokine receptors (Preite et al., 2019). Loss-of-function and gain-of-function mutations of p110δ, encoded by the *PIK3CD* gene, lead to immunodeficiency and immune dysregulation disorders in humans, respectively (Fruman et al., 2017; Lucas et al., 2016). Affected patients present with variable and complex clinical and immunological manifestations due to the pleiotropic effect of PIK3CD in immune cells, many of which are recapitulated, at least in part, in mouse models.

Treg maintain elevated levels of PTEN, a negative regulator of PI3K activity (Bensinger et al., 2004; Zeiser & Negrin, 2008). This balancing activity of PTEN suggests that PI3Kδ modulates a critical signaling program(s) in Treg homeostasis and activity. Consistent with this concept, mice lacking PI3Kδ have reduced numbers of Treg in the periphery and develop colitis, and PI3Kδ-deficient Treg cells fail to protect mice against experimental colitis (Patton et al., 2006). Conversely, PI3Kδ-deficent mice and mice lacking p110δ specifically in Treg cells mount improved protective anti-tumor immune responses targeting a broad range of cancers including solid tumors. In part, this leads to unleashing enhanced effector CD8 T responses within the tumor microenvironment (Ali et al., 2014). Similarly, in a Treg-specific FOXO1 gain-of-function mouse model, the resulting depletion of Treg within in the tumor microenvironment leads to more effective anti-tumor immunity (Luo et al., 2016). In contrast, Treg-specific deletion of PTEN leads to enhanced PI3K activity, resulting in a systemic lymphoproliferative disorder despite an increase in the CD25^-^ subset of Treg (Huynh et al., 2015). Furthermore, consistent with the human disorder, activated PI3Kδ syndrome (APDS), mice expressing activating mutations in PIK3CD display lymphoproliferation and intestinal and systemic inflammation (Preite et al., 2018). In this model, referred to hereafter as activated (a)PIK3CD mice, global expression of aPIK3CD leads to a significant increase in the activated and effector CD4 and CD8 T cells despite an increase in FoxP3^+^ Treg (Bier et al., 2019; Jia et al., 2021; Preite et al., 2018). Interestingly, mixed bone marrow chimeras (established using hematopoietic stem cells derived from aPIK3CD and control mice) revealed that the aberrant differentiation of CD4 T cells resulted predominantly from extrinsic factors driven by alternative cell types (Bier et al., 2019; Preite et al., 2019). Thus, while previous studies have established the significant role of PI3Kδ signaling in Treg homeostasis and function, the specific role for aPIK3CD signaling in differentiation and function of Treg remains incompletely defined.

To better understand the cell-intrinsic function of the PI3Kδ pathway in Treg homeostasis, we generated mice expressing aPIK3CD specifically within Foxp3^+^ expressing cells. Here, we show that Treg specific-aPIK3CD mice exhibit a significant increase in Treg numbers. Surprisingly, in parallel, Treg specific aPIK3CD animals exhibit a progressive increase in memory/effector CD4 and CD8 T cells with an increased production of inflammatory cytokines and decreased proportions of naïve T cells. Consistent with CD4 T cell activation, Treg specific-aPIK3CD animals exhibited increased spontaneous GC responses and produced high-titers of IgG and IgM autoantibodies targeting a broad array of self-antigens. Additional analyses revealed that aPIK3CD dysregulates IFN-γ production in Treg and negatively regulates PD1 expression. Finally, young Treg specific aPIK3CD mice mounted a more robust humoral response upon immunization with a T-cell dependent antigen compared to controls, leading to an increase in the number of antigen specific GC B cells and plasma cells as well as a reduction in the proportion of T follicular regulatory (Tfr) cells.

## Results

### Generation and characterization of Treg-specific aPI3KCD mutant mice

To examine the impact of a gain-of-function mutation in PI3Kδ signaling in Treg development and function, Foxp3^Cre^ mice were crossed with mice harboring a loxP-flanked exon within the endogenous *Pik3cd* gene encoding the murine equivalent of the most common gain-of-function mutation of PIK3CD in APDS patients [E1021K (E1020K in mice] (Al Qureshah et al., 2021; Wray-Dutra et al., 2018). The Foxp3^Cre^ strain expresses Cre recombinase and cis-linked YFP specifically within Treg cells (Rubtsov et al., 2008), and this breeding strategy was designed to promote expression of aPIK3CD only within Treg (*Foxp3*^Cre^ *Pik3cd*^E1020K/+^ hereafter referred as Foxp3-aPIK3CD mice). Only male mice were used for our studies, as heterozygous female mice lacked expression of Cre recombinase in ∼50% of Treg secondary to random X-inactivation (due to location of *Foxp3* gene on the X-chromosome). Treg from Foxp3^Cre^ wild type (hereafter referred as Foxp3-WT) and Foxp3-aPIK3CD mice are marked by YFP (shown schematically in Figure 1A). To confirm specificity of Cre-mediated ‘flipping’ of the LoxP-flanked exon containing the mutant allele in Treg, we purified and analyzed Treg (CD4^+^CD25^+^) from Foxp3-aPIK3CD mice for the presence of the aPIK3CD allele within genomic DNA using a droplet digital (dd)PCR-based assay (Wray-Dutra et al., 2018). As expected, expression of the mutant allele was present with an allele frequency of ∼50%, consistent with heterozygous expression in the majority of Treg (Figure 1B).

**Figure 1.**
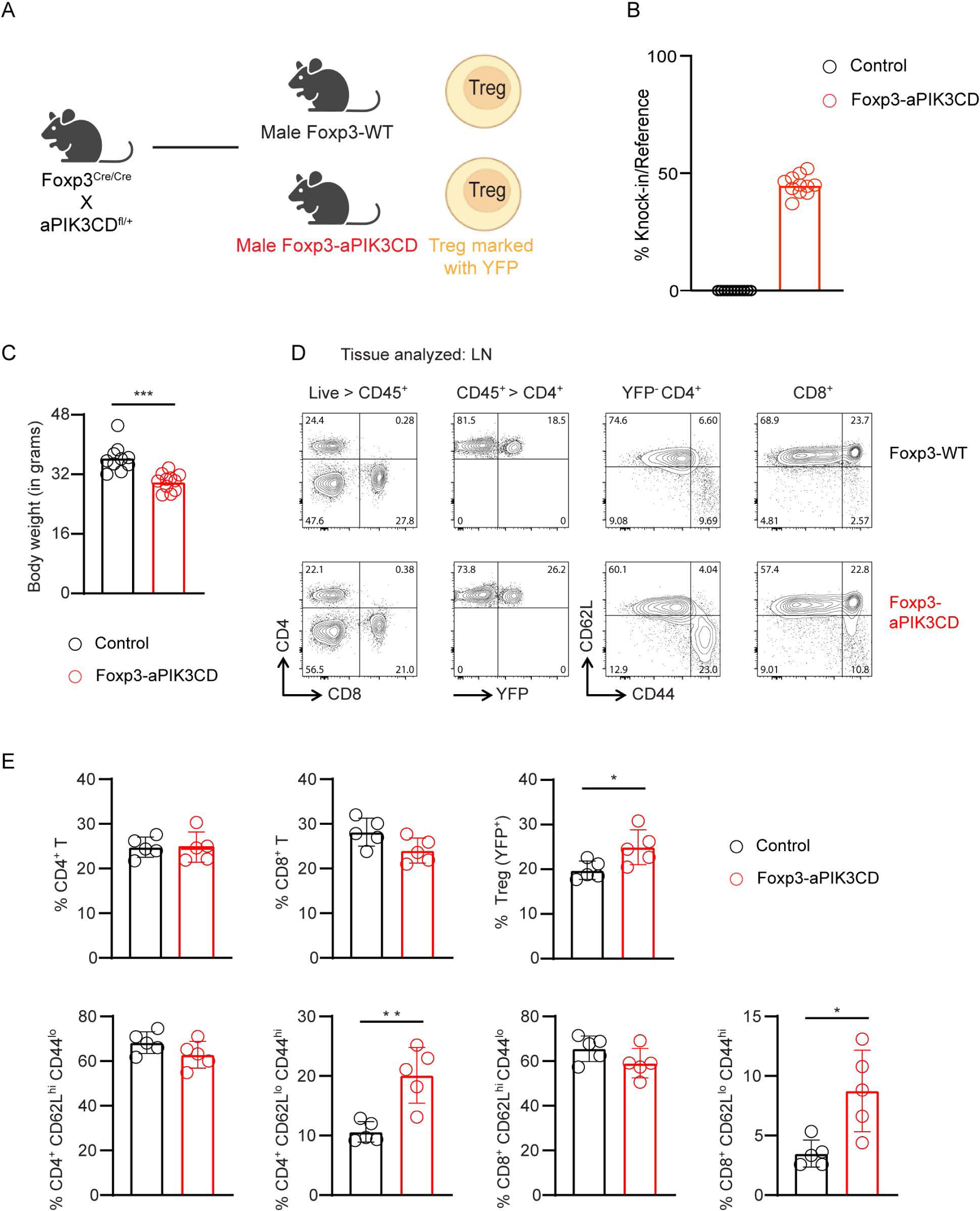
Generation and characterization of Treg-specific aPIK3CD mutant mice. (A) Schematic representation of the generation of Foxp3-WT and Foxp3-aPIK3CD mice by crossing homozygous male aPIK3CD with homozygous female Foxp3^Cre^ mice. (B) ddPCR analysis of genomic DNA isolated from purified Treg (CD4^+^YFP^+^) from the spleens and lymph nodes of Foxp3-aPIK3CD mice. Data is expressed as frequency of knock-in probe binding normalized to reference probe binding. Expected frequency of an aPIK3CD heterozygote mouse is ∼50% in the presence of Cre. Data is combined from two independent experiments; technical replicates were averaged, and each dot represents a biological replicate. (C) Body weight of Foxp3-aPIK3CD and age matched control Foxp3-WT mice at the time of analysis (32-35 weeks of age), n ≥ 10 mice per group. (D) Representative flow plots and (E) relative frequency of T cell subsets in Foxp3-WT and Foxp3-aPIK3CD mice. D, left two panels and E, top panels: Representative flow plots and graphs showing the frequency of CD4^+^, CD8^+^, and Treg (CD4^+^YFP^+^) in LNs. D, right two panels and E, bottom panels: Naïve (CD62LhighCD44low) and effector (CD62LlowCD44high) CD4^+^ and CD8^+^ T cells from LNs. Two independent experiments were performed (n ≥ 3 mice per group). Significance determined by t-test (unpaired). *, P < 0.05, **, P < 0.01. Graphs depict mean.

Foxp3-aPIK3CD mice did not exhibit abnormalities in appearance or weight for up to ∼20 weeks. However, aged Foxp3-aPIK3CD animals (32-35 wks) exhibited a decrease in body weight relative to age and sex matched control animals (Figure 1C); findings potentially suggestive of chronic inflammation. To begin to address this observation, we examined the frequency and activation status of T cells in mice at 32 to 35 weeks of age. We first assessed the frequency of CD4^+^ and CD8^+^ T cells and Treg (CD4^+^YFP^+^) in lymph nodes (LNs). While the frequency of CD4^+^ and CD8^+^ T cells was not altered, the proportion of Treg was significantly higher in Foxp3-aPIK3CD mice (Figure 1D, left panels and 1E, top panels). To gain insight into T cell activation and Treg function, we analyzed CD4^+^T and CD8^+^T cells for naïve/effector phenotypes. Despite the significantly higher frequency of Treg among CD4^+^T cells in Foxp3-aPIK3CD mice, both CD4^+^T and CD8^+^T cells contained significantly higher proportions of CD62L^low^CD44^high^ memory/effector CD4^+^T and CD8^+^ T cells and a trend towards a decrease in CD62L^high^CD44^low^ naïve T cells (Figure 1D, right panels and 1E, bottom panels). Together these observations demonstrate that Treg-specific expression of aPIK3CD leads altered health status based on progressive weight loss and correlates with a higher proportion of total Treg within the periphery and a seemingly paradoxical defect in tolerance leading to activation of both CD4 and CD8 Teff within the lymphoid compartment.

### Spontaneous GC formation and B cell autoimmunity in Foxp3-aPIK3CD mice

Based upon the broad impact of Treg-intrinsic aPIK3CD expression within the T cell compartment, we next examined its potential role in altering B cell function. We first assessed spontaneous germinal center (GC) B cell populations in young (12wk) and aged (32wk) Treg-aPIK3CD and control mice. Young Foxp3-aPIK3CD mice exhibited a trend toward higher proportion of GL7^+^CD38-GC B cells (Supplementary Figure 1A). Strikingly, there was a significant accumulation of GC B cells in older Treg-aPIK3CD mice compared to WT control with an increase in both the proportion and absolute number of GC B cells (Figure 2A, B). We did not observe a difference in the CD138^+^ plasma cells (PC) in young or old mice (Figure 2A, C; Figure 2–figure supplement 1B).

**Figure 2.**
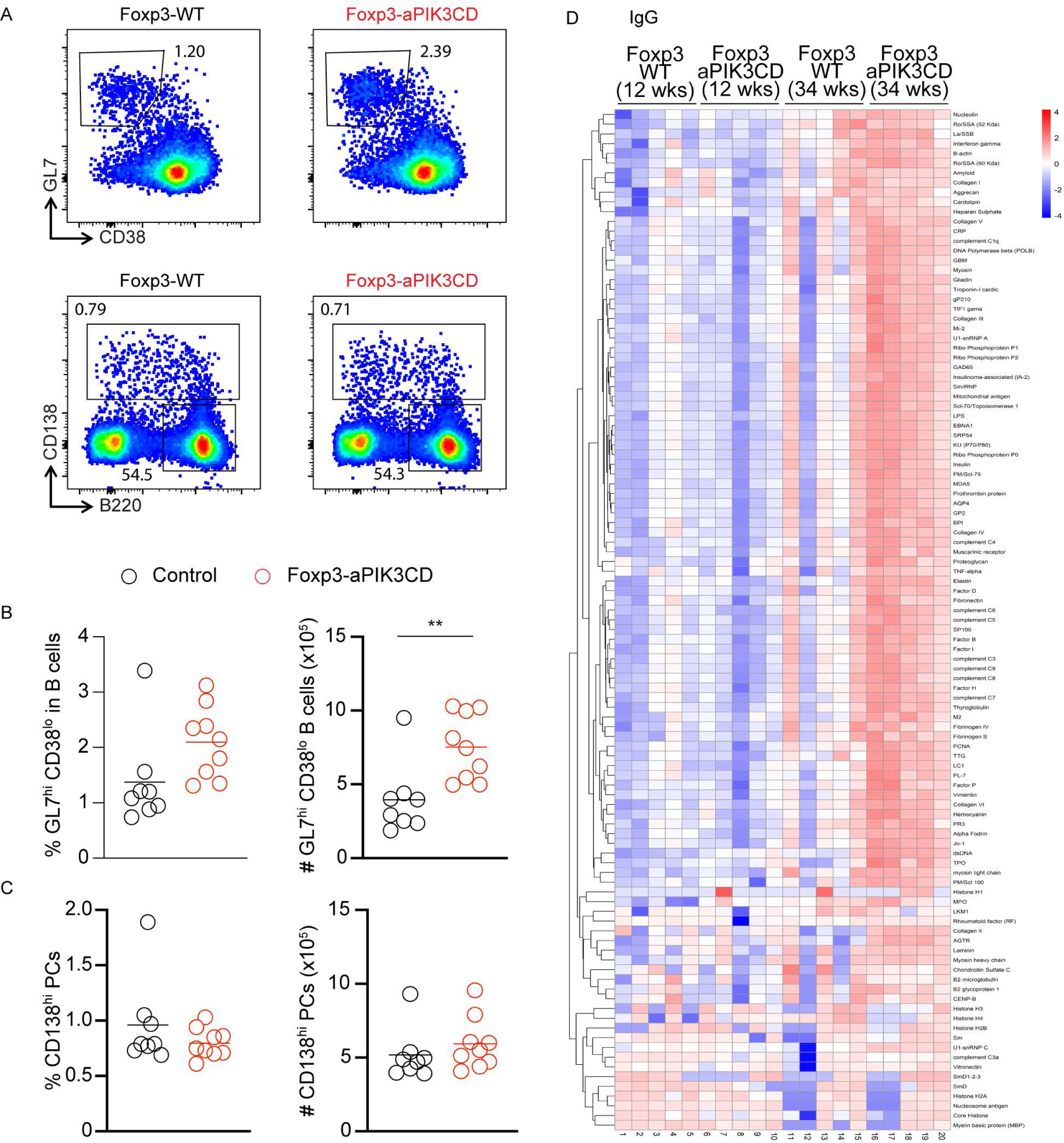
Foxp3-aPI3KCD mice develop spontaneous GC responses and autoantibodies. (A) Representative flow plots showing GC B cells (GL7^+^CD38−) B cells (gated on B220^+^) and plasma cells (CD138^+^) in the spleen of 34-wk-old Foxp3-WT and Foxp3-aPIK3CD mice. (B) Frequency and absolute number of GC B cells in the spleen of 34-wk-old Foxp3-WT and Foxp3-aPIK3CD mice. (C) Frequency and absolute number of CD138^+^ PCs in the spleen of 34-wk-old Foxp3-WT and Foxp3-aPIK3CD mice. (D) Autoantibody microarray heatmap shows signal intensity of IgG autoantibodies to the most significant autoantigens in the serum of young (12wks) and old (34wks) Foxp3-WT and Foxp3-aPIK3CD mice. Two independent experiments were performed (n ≥ 3 mice per group). Significance determined by t-test (unpaired). **, P < 0.01. Graphs depict mean.

The increase in GC B cells in older Foxp3-aPIK3CD mice raised the question of whether Treg-intrinsic aPIK3CD would promote autoantibodies in mice. To address this question, we performed an autoantigen microarray of serum samples from young and old Foxp3-aPIK3CD mice and age matched Foxp3-WT controls. Consistent with the GC data, young Foxp3-aPIK3CD showed a modest increase in IgM reactivity to autoantigens and older Foxp3-aPIK3CD mice exhibited high levels of both IgM and IgG autoantibodies reactive across a wide range of autoantigens (Figure 2D; Figure 2-figure supplement 1C). Together, these data show that Treg-intrinsic aPIK3CD expression promotes the formation of spontaneous GC responses leading to the production of IgM and IgG autoantibodies in older mice.

### Increased thymic Treg precursor development in Foxp3-aPIK3CD mice

To better understand the potential events leading to the higher proportion of Treg in secondary lymphoid tissues in Foxp3-aPIK3CD mice, we next assessed Treg development within the thymus. Comparable frequencies of double positive (DP) and single positive (SP) CD4^+^ and CD8^+^ T cells were present in Foxp3-WT and Foxp3-aPIK3CD mice (Figure 3A, left panels and 3B, first two panels). However, the proportion of Treg precursors (CD4 SP CD25^+^YFP^-^ T cells) was significantly higher in Foxp3-aPIK3CD in comparison to Foxp3-WT mice (Figure 3A, middle panels and 3B, second panel from right). This finding suggested that Foxp3-aPIK3CD mice might generate higher numbers of naïve Treg within the thymus. Consistent with this concept, we observed a trend for a higher proportion of Treg among CD4 SP population in Foxp3-aPIK3CD mice in comparison to Foxp3-WT mice (Figure 3A, right panels and 3B, right panel). To gain insight into the suppressive function of Treg in the presence of aPIK3CD, we performed in vitro suppression assays using Treg isolated from Foxp3-aPIK3CD and Foxp3-WT mice. aPIK3CD and control Treg exhibited comparable suppressive capacity in modulation of Teff proliferation (Figure 3C). Taken together, these findings indicate that aPIK3CD has no major negative impact on thymic Treg precursors or Treg and no impact on ex vivo suppressive activity. While these findings do not fully address the production rate of naïve Treg, they suggest that increased Treg precursor frequency may contribute to the higher proportion of Treg within secondary lymphoid tissues.

**Figure 3.**
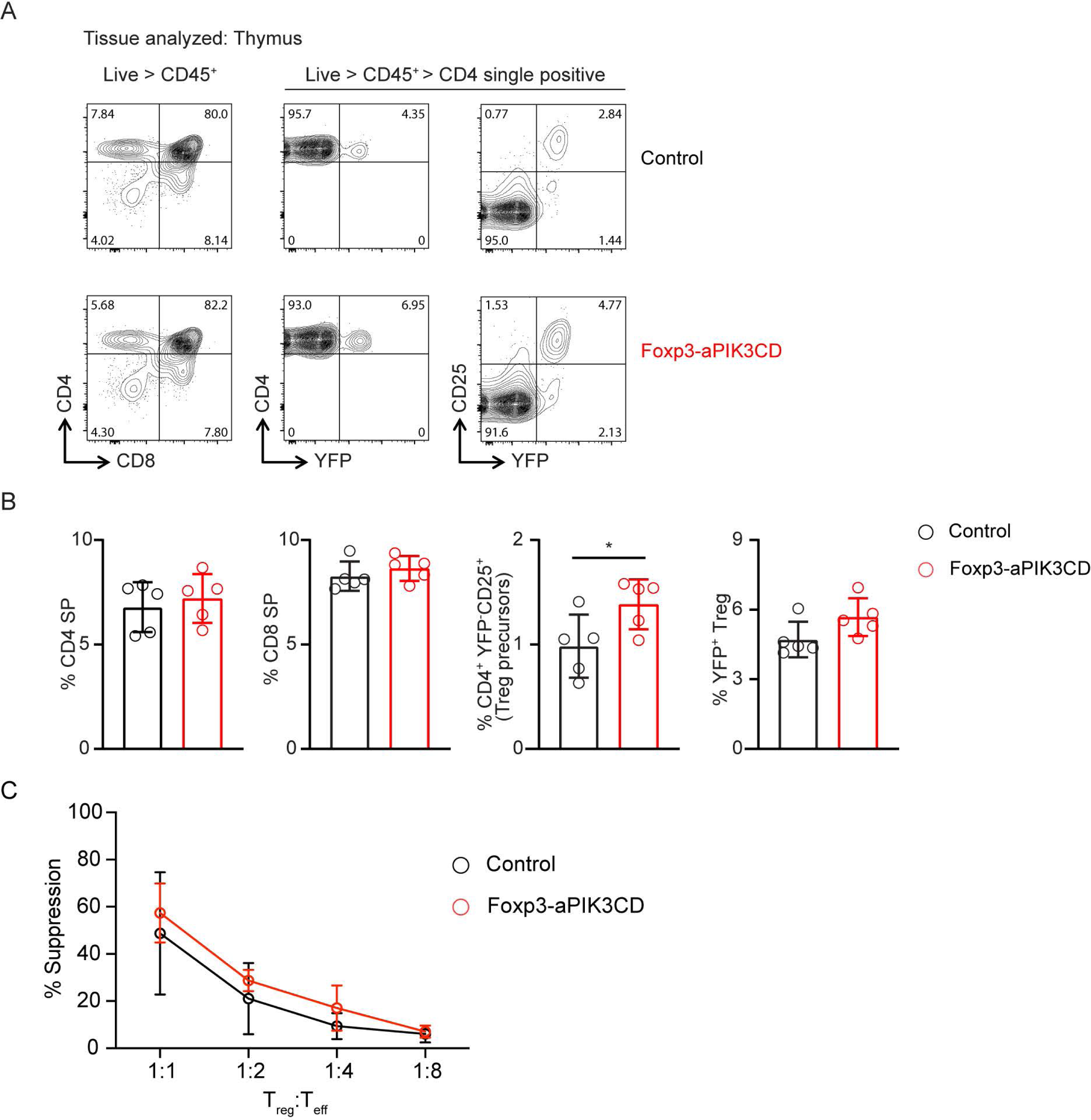
Thymic Treg development and *in vitro* Treg suppressive activity in Foxp3-aPIK3CD mice. (A) Representative flow plots showing the development of CD4 or CD8 single positive (SP) T cells (left panels), Treg precursors (middle panels) and Treg (right panels) in thymus of Foxp3-WT vs. Foxp3-aPIK3CD mice. (B) Graphs showing the frequency of CD4 and CD8 SP T cells (left two panels), Treg precursor (CD4 SP, CD25^+^YFP^-^ T cells) and Treg (CD4^+^YFP^+^ T cells). Two independent experiments were performed (n ≥ 3 mice per group). C. Percent suppression of CellTrace Violet-labeled effector CD4 T cells (Teff) by Treg isolated from Foxp3-aPIK3CD vs. wild-type C57BL/6 mice expressed as a function of the ratio of Treg to Teff. Significance determined by t-test (unpaired). *, P < 0.05. Graphs depict mean.

### Dysregulated production of IFN-γ by Teff and Treg in Foxp3-aPIK3CD mice

To further explore the functional profile of Treg expressing aPIK3CD, we analyzed the production of proinflammatory cytokines IFN-γ and IL-17 in Treg and in both CD4^+^and CD8^+^ Teff in cohorts of 30-34 wk old mice. To our surprise, LN resident Treg expressing aPIK3CD produced significantly elevated levels of IFN-γ. This feature, however, was not observed for Treg in spleen or other peripheral tissues including the lungs (Figure 4A-B, left panels). As Treg are essential for controlling peripheral tolerance by regulating self-reactive Teff, we assessed whether altered Treg cytokine production correlated with dysregulated Teff responses. In Foxp3-aPIK3CD mice, both CD4^+^ and CD8^+^ Teff produced significantly more IFN-γ in LN and spleen (Figure 4A-B, middle and right panels) consistent with a chronic inflammatory response that worsened with age. We did not observe changes in IL-17 production from CD4^+^T cells in the presence of aPIK3CD (data not shown).

**Figure 4.**
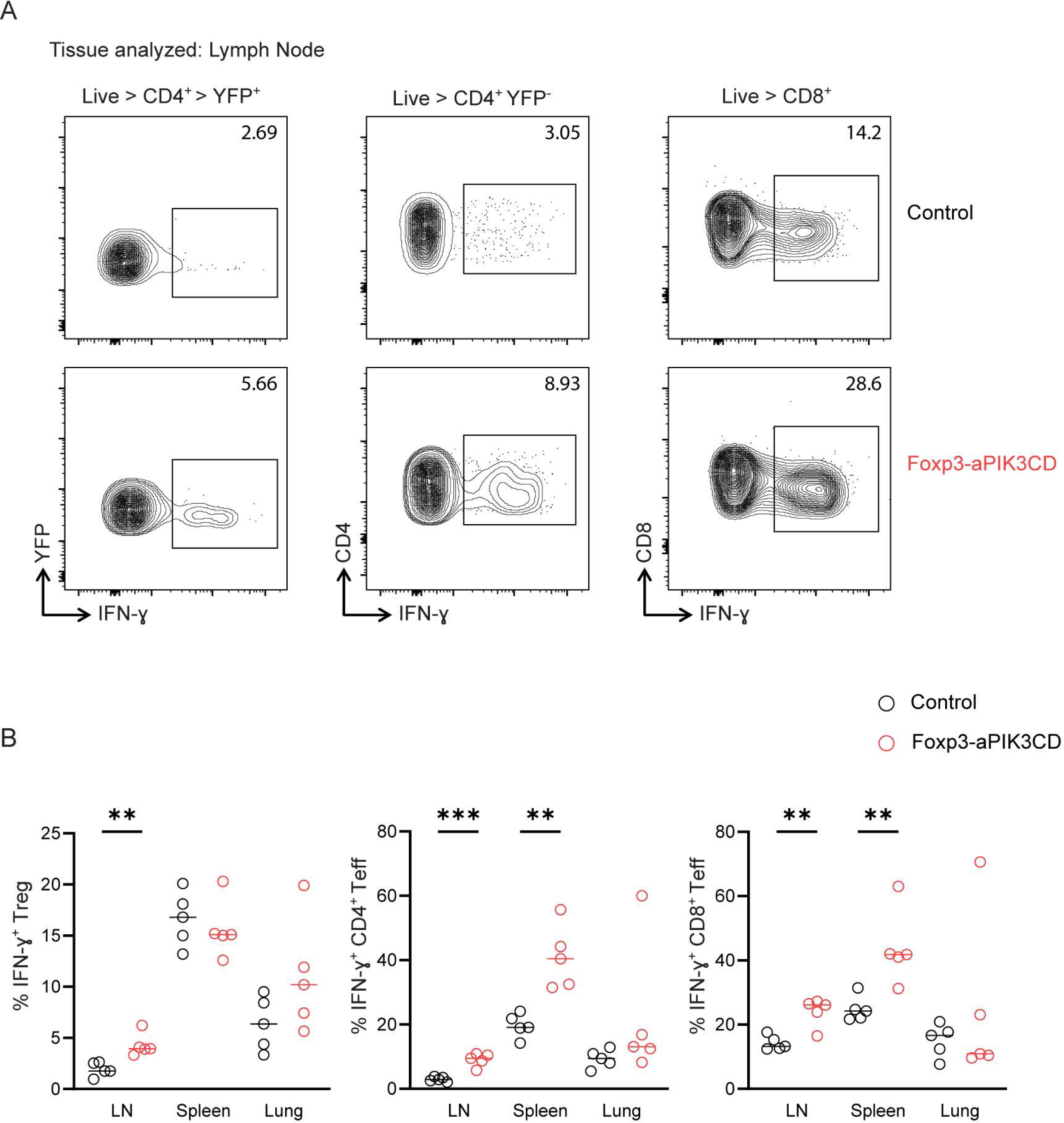
Dysregulated IFN-γ cytokine production by Treg and Teff in Foxp3-aPIK3CD mice. (A) Representative flow plots showing the IFN-γ cytokine production by Treg (left panels), CD4^+^ Teff (middle panels) and CD8^+^ Teff (right panels) isolated from LNs of Foxp3-WT vs. Foxp3-aPI3K3CD mice. (B) Graphs showing the frequency of IFN-γ producing Treg, CD4^+^ and CD8^+^ T cells isolated from LNs, spleen and lung of Foxp3-WT and Foxp3-aPI3K3CD mice. Two independent experiments were performed (n ≥ 3 mice per group; ages 30-34 wks). Significance determined by t-test (unpaired). **, P < 0.01, ***, P < 0.001. Graphs depict mean.

### Treg in Foxp3-aPI3KCD mice exhibit enhanced PD1 expression

Next, we performed detailed Treg immunophenotyping to help determine the potential impact of aPIK3CD expression on Treg function. First, we asked if Treg from Foxp3-aPIK3CD mice had defective expression of the transcription factor Foxp3, the master transcriptional regulator of Treg. We observed no significant impact on Foxp3 expression in Treg isolated from the thymus, spleen, or lungs of Foxp3-aPIK3CD mice (Figure 5–figure supplement 2A). Slightly higher levels of Foxp3 expression were observed in Treg from LNs of Foxp3-aPIK3CD mice in comparison to Foxp3-WT mice (Figure 5–figure supplement 2A, top left panel). Next, we assessed the expression of CD25, the alpha subunit which confers high affinity to the IL2R complex and is constitutively expressed on the surface of Treg. We observed no defect in the frequency of CD25^+^ Treg analyzed from indicated tissues (Figure 5A, top panels and Figure 5B, top left panel) and no reduction in CD25 cell surface expression levels (Figure 5–figure supplement 2B, top right panel), suggesting that the impact of aPIK3CD expression is not attributable to defects in Foxp3 and/or CD25 expression.

**Figure 5.**
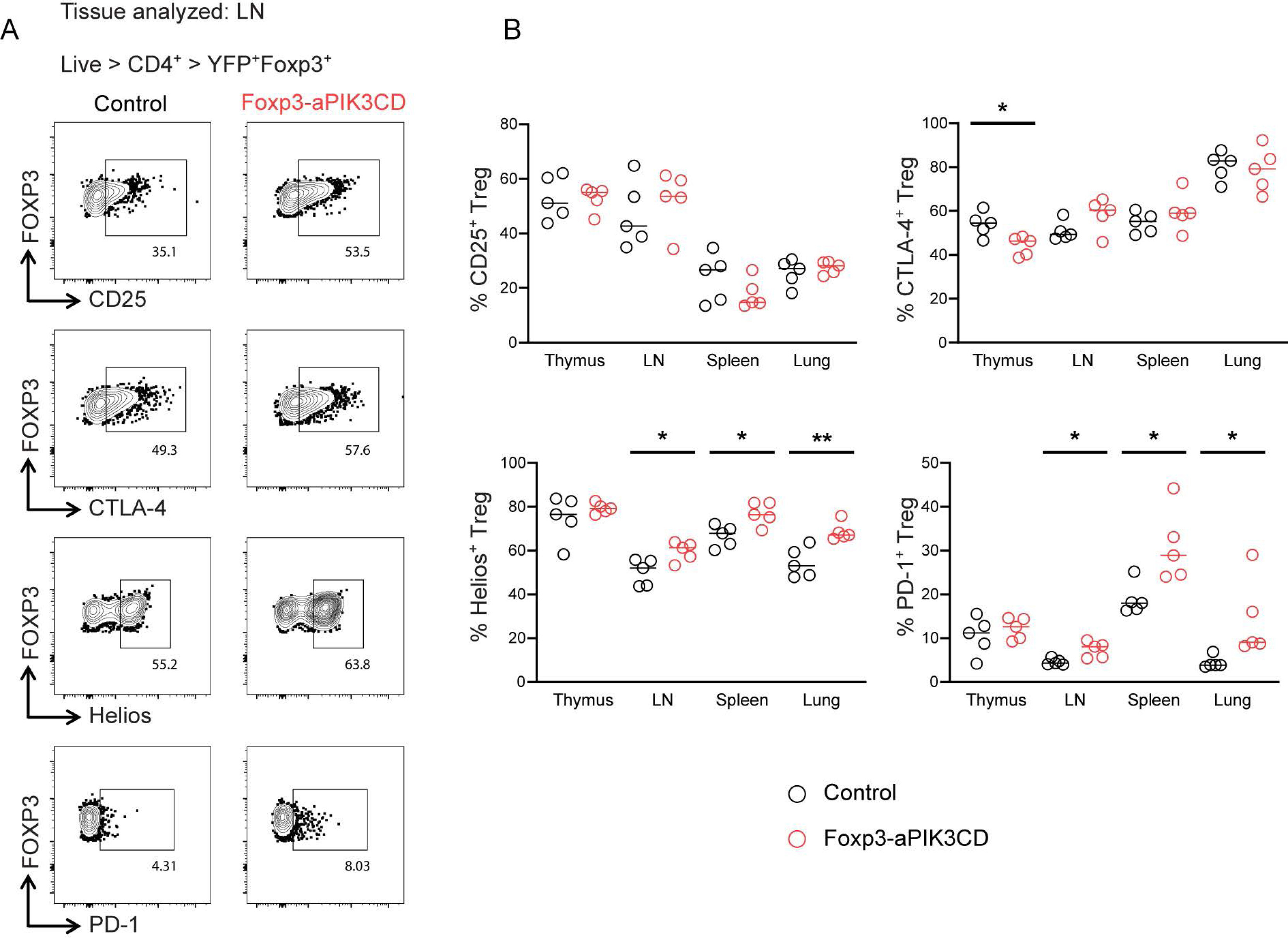
Treg expressing aPIK3CD exhibit phenotypic changes including increased PD-1 levels. (A) Representative flow plots showing the expression of Treg associated proteins in Treg isolated from LNs of Foxp3-WT (left panels) and Foxp3-aPIK3CD mice (right panels). (B) Graphs showing the frequency of Treg expressing indicated proteins (CD25, CTLA-4, Helios, PD-1) in thymus, LNs, spleen and lung of Foxp3-WT and Foxp3-aPIK3CD mice. Two independent experiments were performed (n ≥ 3 mice per group; ages 30-34wks). Significance determined by t-test (unpaired). *, P < 0.05 **, P < 0.01, Graphs depict mean.

CTLA-4 expression on Treg is essential for effective regulation of naïve T cell activation and in the prevention of autoimmunity (Jain et al., 2010). Thus, we assessed the expression of CTLA-4 on Treg from Foxp3-aPIK3CD mice and compared the frequency and expression levels with Treg from Foxp3-WT mice. Consistent with a lack of defects in Foxp3 and/or CD25 expression, we observed no change in the frequency of CTLA-4^+^ Treg or in the expression level of CTLA-4 on peripheral Treg from Foxp3-aPIK3CD mice (Figure 5A second panel from top and Figure 5B, right upper panel). However, we did observe a modest decrease in the frequency of CTLA-4^+^ Treg in the thymus of Foxp3-aPIK3CD mice (Figure 5B, right upper panel).

Next, to address whether the increased frequency of Treg in secondary lymphoid tissue of Foxp3-aPIK3CD mice reflected increased egress from the thymus vs. increased generation of peripheral Treg, we analyzed Treg based expression of the transcription factor, Helios. As Helios expression in mice is restricted to thymus-derived Treg (Thornton et al., 2010), we reasoned that an increased proportion of Helios^+^ Treg may indicate more efficient production and/or egress of Treg from the thymus. Consistent with the increased frequency of thymic Treg precursors and the increased proportion of total YFP^+^ Treg in Foxp3-aPIK3CD mice, the proportion of Helios^+^ Treg was increased in peripheral tissues including LNs and spleen in mutant mice (Figure 5A, third panels from top and Fig 5B, bottom left panel). Thymic Treg express Helios whereas peripheral Treg are comprised of both Helios^+^ and Helios^-^ populations in adult mice (Thornton et al., 2010; Thornton et al., 2019). As anticipated, based on the above data, we observed a similar frequency of Helios^+^ cells within gated YFP^+^ Treg in the thymus in Foxp3-aPIK3CD vs. control mice and Helios^+^ cells exhibited similar MFI across strains in all the tissues (Figure 5B, bottom left panel –figure supplement 2A). Together, these data suggest that increased thymic production leads to an increased proportion of thymus-derived Helios^+^ Treg in Foxp3-aPIK3CD mice.

Recent studies have revealed a critical and non-redundant role for PD-1 in Treg suppressive function (Kamada et al., 2019; Tan et al., 2021). Notably, PD-1 negatively regulates Treg function via enhanced PI3K and AKT dependent signaling. Therefore, despite the increased proportion of Treg, we asked whether Foxp3-aPIK3CD mice had an increased frequency of PD-1^+^ Treg; a finding that might result in Treg with reduced suppressive function thereby promoting inflammation in older Foxp3-aPIK3CD mice. We examined the frequency of PD-1^+^ Treg and the expression level of PD-1 on Treg. Strikingly, Foxp3-aPIK3CD mice had significantly higher proportion of PD-1^+^ Treg in all the tissues examined except the thymus (Figure 5A, bottom panels and 5B, bottom right panel). Interestingly, Treg in Foxp3-aPIK3CD mice also exhibited a trend for increased PD–1 expression based upon MFI of antibody staining (Figure 5–figure supplement 2B, middle right panel). Together, these data support the concept that aPIK3CD negatively regulates Treg function by promoting increased PD-1 expression and engagement.

### Treg demonstrate reduced competitive fitness in the presence of aPI3KCD expression

The above findings failed to identify a deficit in Treg development in the presence of aPIK3CD. Despite this, we observed a significant increase in both frequency of Treg precursors in the thymus and total Treg within the secondary lymphoid tissues. Thus, based on these findings, we next sought to determine whether aPIK3CD Treg might exhibit altered competition in the presence of WT Treg. To address this question, we used Foxp3^Cre/WT^ aPIK3CD heterozygous female mice and control (Foxp3^Cre/WT^ WT) mice as depicted (Figure 6A). In the former strain, due to random inactivation of X chromosome, both WT and aPIK3CD Treg are generated in the same animal. This strain is anticipated to be healthy and have a normal life span due to the presence of both WT and aPIK3CD Treg. Consistent with this hypothesis, Foxp3^Cre/WT^ aPIK3CD heterozygous females showed no signs of disease or weight loss when evaluated at ∼35 weeks of age.

**Figure 6.**
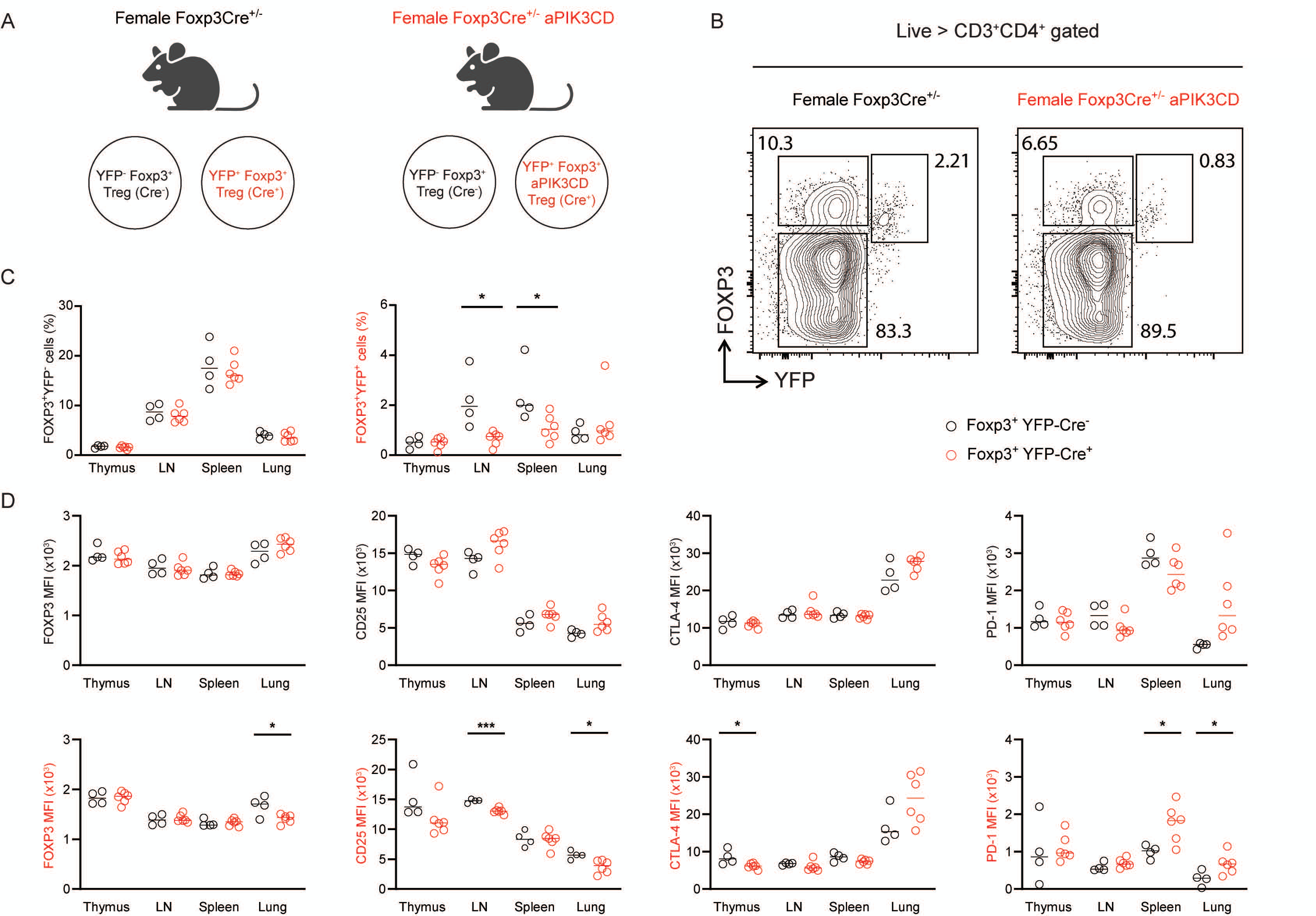
Treg expressing aPIK3CD exhibit a competitive disadvantage in vivo. (A) Schema showing the strategy for tracking WT vs aPIK3CD Treg in vivo in heterozygous carrier female mice. (B) Representative flow plots used to identify Foxp3^+^YFP^-^ (upper left gate) and Foxp3^+^YFP^+^ (upper right gate) in female Foxp3^Cre/WT^ control (left) vs. Foxp3^Cre/WT^ aPIK3CD (right) “test” mice. (C) Graphs showing the frequency of control Foxp3^+^YFP^-^ (left) and test Foxp3^+^YFP^+^ (right) Treg in thymus, LNs, spleen and lung of Foxp3^Cre/WT^ vs. Foxp3^Cre/WT^ aPIK3CD female mice. (D) Graphs showing the mean fluorescent intensity of Foxp3, CD25, CTLA-4 and PD-1 on Foxp3^+^YFP^-^ gated Treg (upper panels) and Foxp3^+^YFP^+^ gated Treg (lower panels). Two independent experiments were performed (n ≥ 4 mice per group). Significance determined by t-test (unpaired). *, P < 0.05 ***, P < 0.001, Graphs depict mean.

We analyzed Live>CD3^+^CD4^+^ gated cells for Foxp3 and YFP from the thymus, spleen, LNs and lungs in cohorts of female heterozygous mice at 30-32 weeks of age. When comparing Foxp3^+^YFP^-^ Treg across strains, no differences were observed in frequency or MFI of Foxp3, CD25, CTLA-4 and PD-1 (Figure 6B, 6C, left panel and 6D). Strikingly, the frequency of aPIK3CD expressing Treg (based on gating of Foxp3^+^YFP^+^ in control vs. mutant allele expressing female mice) was significantly decreased in LNs and spleen despite comprising a comparable proportion of cells within the thymus (Figure 6B and C, right panel).

These findings suggested that Foxp3^+^YFP^+^ Treg exhibit a selective disadvantage in the presence of aPIK3CD expression. Consistent with this selective deficit, we observed phenotypic differences when comparing WT vs. aPIK3CD Treg. The expression level of CD25 was reduced in aPIK3CD Treg from LNs and lung, and lower levels of Foxp3 expression were observed in aPIK3CD Treg in the lung (Figure 6D, panels at left). In addition, we noted a reduction in CTLA-4 MFI on thymic aPIK3CD Treg (Figure 6D, second from right). As described above, Foxp3-aPIK3CD Treg exhibit a significantly higher proportion of PD-1^+^ Treg (Figure 5). Notably, consistent with that observation, aPIK3CD Treg from spleen and lung of Foxp3^Cre/WT^ aPIK3CD heterozygous female mice exhibited higher expression levels of PD-1 (Figure 6E, far right panel). As these findings in the competitive setting were consistent with those in aged Foxp3-aPIK3CD mice, they suggest a shared mechanism leading to altered Treg function. Together, these data demonstrate that Treg expressing aPIK3CD exhibit a competitive disadvantage in the presence of WT Treg and that this deficit correlates with increased expression of PD-1.

### Altered T dependent immune responses in Foxp3-aPIK3CD mice

Based upon the altered Treg phenotype and spontaneous GC responses and autoantibodies in aged Foxp3-aPIK3CD animals, we next investigated the impact of aPIK3CD on humoral responses following TD immunization in cohorts of 12- to 14-week-old animals. We immunized Foxp3-WT and Foxp3-aPIK3CD mice with virus-like particles (VLPs) containing a TLR ligand (VLP-ssRNA). At day 14 post immunization, Foxp3-aPIK3CD mice showed an increased proportion and number of GL7^+^CD38-GC B cells compared with controls (Figure 7A). We next assessed VLP-specific B cells within the GC compartments. While the overall frequency of VLP-specific GC was lower, Foxp3-aPIK3CD mice showed significantly increased numbers of VLP-specific GC B cells compared with Foxp3-WT (Figure 7B). Additionally, Foxp3-aPIK3CD mice exhibited enhanced proportion and number of IgG2c^+^ VLP-specific GC B cells (Figure 7C). In parallel, we evaluated the impact of aPIK3CD on PC generation. Consistent with the enhanced GC response, Foxp3-aPIK3CD mice exhibited an increased proportion and number total CD138^+^ PCs, VLP-specific PCs, and IgG2c VLP-specific PCs (Figure 7D-F). Consistent with the flow cytometry data, serum IgG and IgG2c VLP-specific antibody levels in Foxp3-aPIK3CD were slightly increased compared with controls (Figure 7–figure supplement 3).

**Figure 7.**
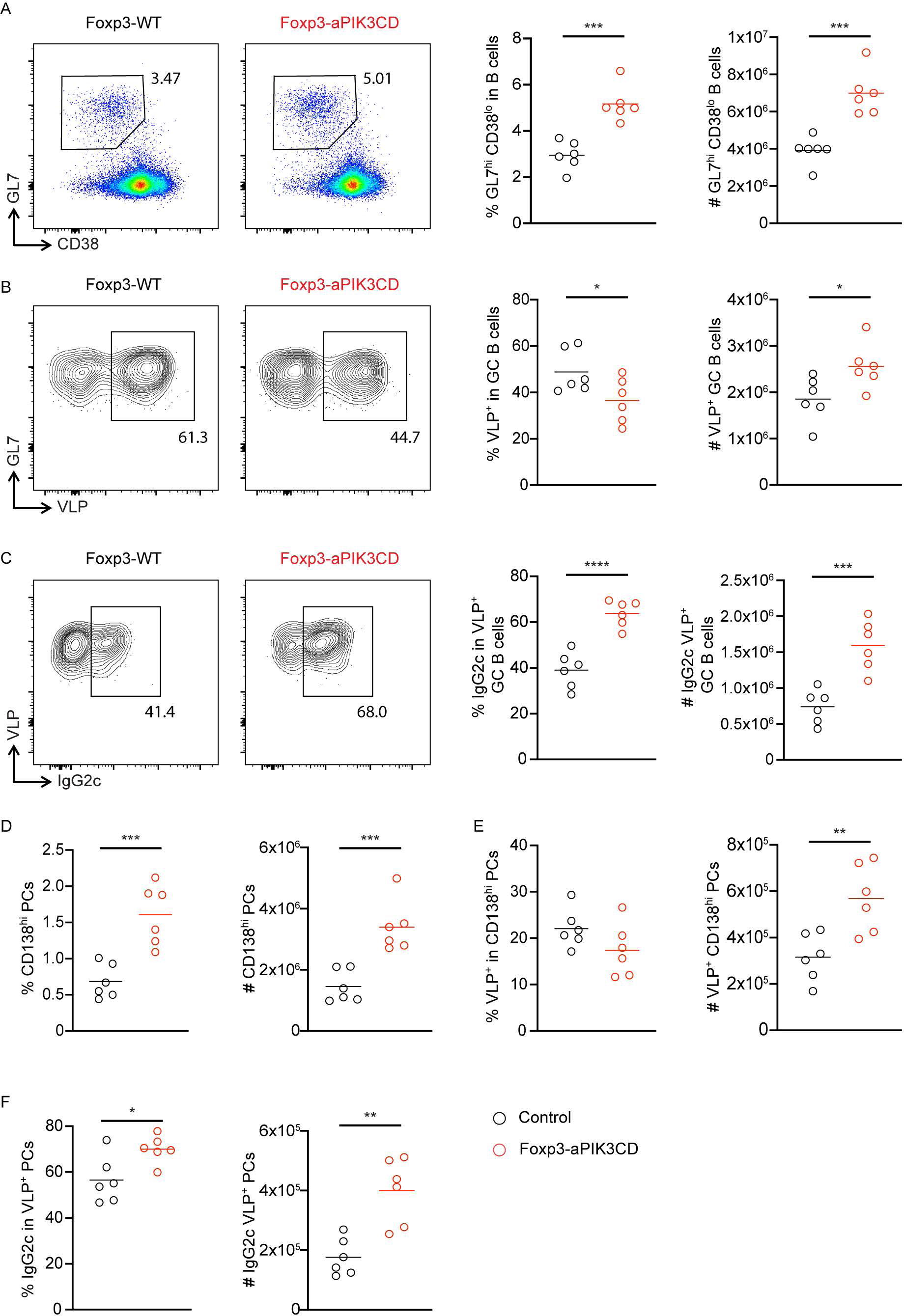
Treg-specific aPIK3CD mice exhibit an enhanced GC response following VLP immunization. (A) *Left plots,* representative flow cytometry analysis of GL7^+^CD38^−^ GC B cells (initially gated on B220^+^ cells) in the spleen of 12- to 14-wk-old Foxp3-WT and Foxp3-aPIK3CD mice 14 days after intraperitoneal immunization with VLP-ssRNA. *Right graphs*, frequencies and absolute numbers of GC B cells. (B) *Left plots,* representative flow cytometry analysis of VLP-specific B cells within GC in mice as described in A. *Right graphs,* frequency and absolute numbers of VLP-specific GC B cells. (C) *Left plots,* representative flow cytometry analysis of IgG2c^+^ VLP^+^ GC B cells. *Right graphs,* frequency and absolute numbers of IgG2c^+^ VLP^+^ GC B cells. (D) Frequency and absolute number of total CD138^+^ PCs in Foxp3-WT or Foxp3-aPIK3CD mice at day 14 after immunization with VLP-ssRNA. (E) Frequency and absolute number of VLP-specific PCs in Foxp3-WT or Foxp3-aPIK3CD. (F) Frequency and absolute numbers of IgG2c^+^ VLP^+^ PCs. Data are combined from two independent experiments (n = 6). Significance determined by unpaired Student’s t test. *, P < 0.05; **, P < 0.01; ***, P < 0.001; ****, P < 0.0001. Graphs depict mean.

Based upon this evidence for an increased humoral response, we hypothesized that Treg intrinsic aPIK3CD expression modulated the function of Tfr cells ultimately leading to altered activity of GC resident T follicular helper (Tfh) following VLP immunization. Consistent with this idea, Foxp3-aPIK3CD mice exhibited a marked increase in the proportion and numbers of Tfh cells (defined as either CD4^+^PD1^+^CXCR5^+^ or CD4^+^PD1^+^ICOS^+^ T cells) in the spleen at day15 post-immunization (Figure 8A-B). In contrast, despite the general increase of Treg observed in Foxp3-aPIK3CD mice, Foxp3-aPIK3CD mice showed a reduced proportion of Tfr (YFP^+^) and a markedly increased Tfh:Tfr ratio compared to controls (Figure 8C). Together, these data show that Foxp3-aPIK3CD mice exhibit an increased humoral response upon immunization with a TD antigen that correlates with a reduction in the proportion of Tfr. These data demonstrate that activated PI3Kδ signaling within Treg leads to impaired regulation of GC B cell responses, likely due to its impact on the functional stability of Tfr cells.

**Figure 8.**
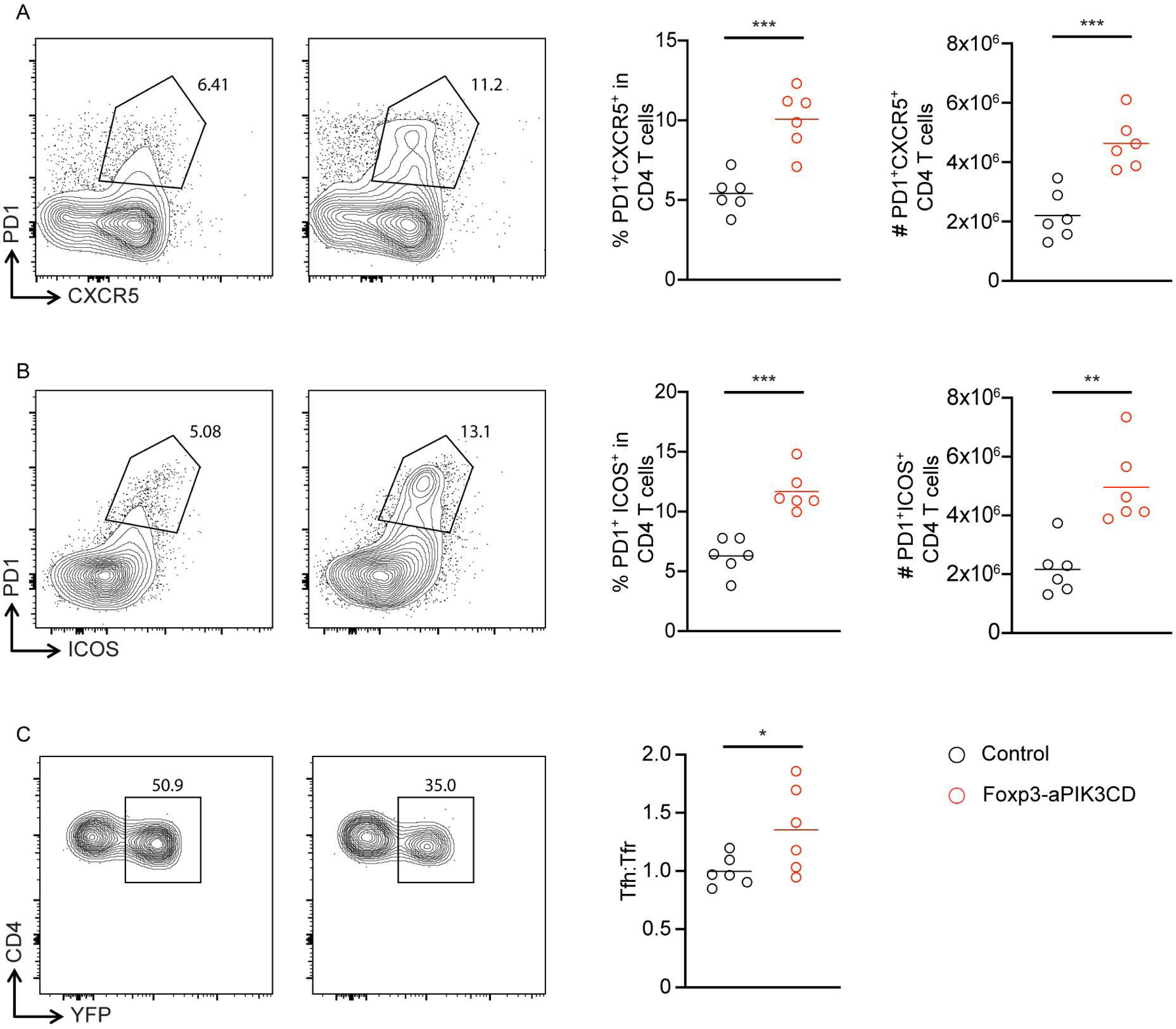
Treg-specific aPIK3CD mice exhibit a skewed Tfh:Tfr ratio following VLP immunization. (A) Left plots show representative flow cytometry analysis of PD1^+^CXCR5^+^ Tfh cells (gated on CD4^+^ cells) in the spleen of 12- to 14-wk-old Foxp3-WT and Foxp3-aPIK3CD mice at day 14 d after intraperitoneal immunization with VLP-ssRNA. Right graphs show frequency and absolute numbers of PD1^+^CXCR5^+^ Tfh cells. (B) Left plots show representative flow cytometry analysis of PD1^+^ICOS^+^ Tfh cells (gated on CD4^+^ cells) in the spleen of 12- to 14-wk-old Foxp3-WT and Foxp3-aPIK3CD mice at day 14 after immunization with VLP-ssRNA. Right graphs show frequency and absolute numbers of PD1^+^ICOS^+^ Tfh cells. (C) Left plots show representative flow cytometry analysis of Tfr cells gated on YFP^+^ cells within the PD1^+^CXCR5^+^ T cell population. Right graphs show ratio of Tfh (YFP^-^):Tfr (YFP^+^) cells. Data are combined from two independent experiments (n = 6). Significance determined by unpaired Student’s t test. **, P < 0.01; ***, P < 0.001. Graphs depict mean.

## Discussion

PIK3CD signaling plays a crucial role in immune homeostasis. Activating mutations in PIK3CD lead to the complex primary immunoregulatory disorder APDS (Lucas et al., 2016), impacting multiple immune cell subsets and manifesting in features of both immune deficiency and immune dysregulation (Coulter et al., 2017; Hartman et al., 2015). As altered Treg function may contribute to these phenotypes, previous studies evaluated Treg frequency in APDS patients based on FOXP3, CD25, and/or CD127 expression with variable results ranging from low, normal, or high Treg:Teff frequencies (Bier et al., 2019; Kotwal et al., 1986; Thauland et al., 2019). Here, we directly assessed the cell intrinsic role for aPIK3CD in Treg development, maintenance/fitness, and function using a conditional KI mouse model. While Treg-specific aPIK3CD expression had minimal impact on thymic Treg development, aPIK3CD expression led to increased numbers of peripheral Treg. Strikingly, aged cohorts of Treg-specific aPIK3CD mice exhibited progressive weight loss, immune dysregulation, and chronic inflammation by ∼30 weeks of age. These findings correlated with increased frequencies of Teff (CD4 and CD8), IFN-γ production, GC formation, autoantibody production and altered Treg phenotypes. Most notably, aPIK3CD Treg exhibited an increase in PD-1 expression, a feature previously correlated with reduced Treg suppressive function (Kamada et al., 2019; Lowther et al., 2016; Tan et al., 2021). Consistent with altered regulation of GC T:B cell interactions, young aPIK3CD mice manifested an enhanced response to immunization with a TD antigen, a finding that correlated with a reduced Tfr:Tfh ratio. Finally, consistent with altered fitness, Treg expressing aPIK3CD exhibited a competitive disadvantage in the presence of WT Treg under homeostatic conditions.

Prior work has demonstrated a requirement for PIK3CD in Treg suppressive function and peripheral maintenance. Mice lacking catalytically active PIK3CD exhibit reduced peripheral Treg suppressive function, reduced Treg frequency, and an altered Treg phenotype despite a ∼2-fold increase in thymic Treg (Patton et al., 2006; Stark et al., 2020). In addition, Treg specific PIK3CD deletion led to reduced growth of B16 melanoma cells mediated by unleashing of CD8^+^ Teff responses (Ali et al., 2014). Interestingly, these animals did not develop spontaneous autoimmunity (Ali et al., 2014; Stark et al., 2020), in contrast to mice with catalytically inactive PIK3CD in all lineages (Patton et al., 2006). In previous work modeling global expression of aPIK3CD (E1020K; identical to the mutation described here), KI mice exhibited an age dependent increase in the frequency of Treg, Teff and Tfh (Bier et al., 2019). Notably, mixed BM chimera studies (using KI and control donor bone marrow) demonstrated that both Teff and Tfh expansion were primarily the result of cell extrinsic factors (Bier et al., 2019; Preite et al., 2019). Thus, it is not clear from previous work how Treg specific aPIK3CD expression might influence Treg development, fitness, or function. In addition, it has not previously been feasible to assess the impact of aPIK3CD on Teff and Tfh cells independently of its direct impacts on Treg mediated function. To address the impact of aPIK3CD on Treg development, we first evaluated Treg precursor and Treg frequencies in the thymus of aPIK3CD mice. aPIK3CD expression led to an increase in Treg precursors and trend for an increase Treg within the thymus. This result was surprising based on previous data showing increased thymic Treg numbers in PIK3CD KO mice suggesting that PI3K signaling negatively regulates thymic Treg development (Patton et al., 2006). In contrast to loss of PI3KCD, defective Treg development was observed following CD4^+^ T cell specific deletion of PDK1, required for downstream PI3K signaling (Park et al., 2010). However, as none of these previous studies were conducted in a cell intrinsic manner, the impact on thymic Treg development may have been indirect. Our data demonstrate that aPIK3CD modestly promotes thymic Treg precursor and Treg numbers and suggest that APDS subjects are likely to exhibit relatively normal Treg thymic development.

Earlier studies, using mouse models of both global (Patton et al., 2006) and Treg intrinsic PIK3CD loss of function (Ali et al., 2014), demonstrate an essential role for PIK3CD signaling in Treg suppressive function. Thus, Foxp3-aPIK3CD Treg animals might have been expected to remain healthy due to increased Treg activity. In contrast to this prediction, aged Foxp3-aPIK3CD mice developed chronic inflammation and autoimmunity, suggesting suboptimal Treg function. Notably, as these animals exhibited an increase in Treg numbers, this process could not be explained by an altered proportion of Treg to CD4^+^ and CD8^+^ Teff. In parallel, we observed no defects in the suppressive function of aPIK3CD Treg in vitro. However, as Treg utilize multiple regulatory mechanisms in vivo to modulate inflammation and tissue repair (Campbell & Rudensky, 2020; Vignali et al., 2008), alterations in these functions may not be reflected in a short-term in vitro suppression assay.

PI3K signaling promotes AKT phosphorylation and its downstream signaling (Manning & Toker, 2017) leading, in part, to phosphorylation and nuclear exclusion of Foxo1 limiting its transcriptional activity (Brown & Webb, 2018). Treg intrinsic loss of Foxo1 results in a scurfy-like disorder and a Treg-intrinsic increase in IFN-γ production—alterations that occur despite normal thymic Treg generation, increased numbers of peripheral Treg, and intact in vitro suppressive function (Ouyang et al., 2012). Our findings in Foxp3^Cre^aPIK3CD mice partially mimic Foxp3^Cre^Foxo1^fl/fl^ mice including increased Treg frequency, IFN-γ production by Treg, and progressive inflammation in older mice. In contrast to complete loss of Foxo1, the more modest inflammatory phenotype in Foxp3^Cre^ aPIK3CD mice likely reflects less robust negative regulation of Foxo1 (a process that remains dependent of downstream aPIK3CD-dependent signals). Additional biochemical studies are required to directly assess the role for aPIK3CD in modulating Foxo1 nuclear exclusion.

PI3K signaling has multiple pleiotropic and developmental-stage dependent impacts across its downstream signaling activities. The altered Treg function in Foxp3^Cre^ aPIK3CD mice and within the competitive setting in Foxp3^Cre/WT^ aPIK3CD heterozygous female mice likely reflects dysregulation of several downstream pathways. Importantly, we observed an increase in PD-1 expression on aPIK3CD-expressing Treg in both settings; a feature predicted to contribute to defective Treg function and chronic inflammation in older mice. PD-1 expression has been shown to inversely impact Treg suppressive function both in humans (Lowther et al., 2016) and mice (Kamada et al., 2019; Tan et al., 2021). PD-1 KO Treg demonstrated superior suppressive function in disease models (EAE and T1D) and exhibited an activated phenotype. Conversely, PD-1 sufficient Treg exhibit a competitive disadvantage in the presence of PD-1 deficient Treg (Tan et al., 2021). Further, loss of AKT or PTEN leads to superior Treg suppressive function in the absence of PD-1 (Crellin et al., 2007; Gerriets et al., 2016; Tan et al., 2021). Similarly, aPIK3CD Treg exhibited a competitive disadvantage (in the presence of WT Treg) that correlated with increased PD-1 expression. In contrast to the progressive disadvantage for aPIK3CD Treg in heterozygous female mice in the setting of immune homeostasis, the increased number of aPIK3CD Treg in Foxp3^Cre^ aPIK3CD mice is likely secondary to inflammation-driven, IL-2 mediated survival and expansion. Based upon the observed chronic inflammation, Teff and Tfh expansion, and autoantibody production in aged Foxp3^Cre^ aPIK3CD, the abundant PD-1^+^ Treg in this setting presumably represent dysfunctional cells. Further, consistent with concept that aPIK3CD impacts multiple pathways, the overall phenotype likely reflects a combination of both a reduced suppressive function due to increased pAKT/Foxo1 and an increased proportion of dysfunctional PD-1^+^ Treg.

PIK3CD plays a crucial role in Tfh activation and GC function including the capacity to support activation, selection and somatic hypermutation (SHM) in GC B cells (Rolf et al., 2010). Further, dysregulated Tfh leads to the development of autoimmunity (Crotty, 2019; Ma & Deenick, 2014; Ueno et al., 2015). As noted above, APDS patients and mice globally expressing aPIK3CD exhibit increased Tfh numbers and altered ratio of Tfh/Tfr (Bier et al., 2019; Preite et al., 2018). However, it remained unknown how aPIK3CD directly impacts Tfr numbers or function. Here, we demonstrate that following VLP immunization in young mice, Foxp3^Cre^ aPIK3CD exhibit increased GC responses characterized by increased numbers of Ag-specific GC B cells, plasma cells and an increase in VLP specific IgG2c—changes that correlated with an enhanced Tfh response as demonstrated by increased numbers of PD1^+^CXCR5^+^ and PD1^+^ICOS^+^ Tfh cells and an increase in the Tfh/Tfr ratio. Together, these findings imply that aPIK3CD leads to a Tfr intrinsic defect that is sufficient to enhance Tfh driven Ab responses. This deficit likely contributes to the development of spontaneous GC responses and broad spectrum of autoantibodies observed in aged Foxp3^Cre^ aPIK3CD animals. Our combined findings, together with previous work suggesting that the Tfh expansion is largely cell extrinsic, imply that a Tfr deficit is likely a significant contributor to the marked Tfh expansion, immune dysregulation, and autoantibodies observed in APDS subjects. Based on the broad array of autoantibodies observed in aged Foxp3^Cre^ aPIK3CD mice, it will be interesting to determine whether the Treg deficits described here also contribute to the enhanced response to commensal bacteria reported in previous studies of global aPIK3CD KI mice (Preite et al., 2018).

In summary, our study emphasizes the important role of PI3Kδ in the development and function of Treg. We found that enhanced PI3Kδ signaling in Treg promotes development yet restricts function leading to autoimmunity that worsened with age. Our data suggests that therapeutic use of PI3Kδ inhibitors may help to prevent or limit systemic autoimmunity in APDS patients and potentially in other autoimmune diseases. Conversely, enhancing PI3Kδ might be useful to perturb Treg cells in cancers allowing for a more effective anti-tumor immune response.

## Materials and Methods

### Mice

Foxp3^YFPCre^ (016959) and B6 (C57BL/6J, 000664) mice were originally purchased from Jackson Laboratory and bred/maintained in specific-pathogen-free facility at Seattle Children’s Research Institute. Pik3cd-E1020K/^+^ mice were generated in our laboratory as previously described (Wray-Dutra et al., 2018). All mice were maintained on C57BL/6 background. Sex and age of mice used in the experiments were indicated in the text. Animal experiments were conducted according to the protocols approved by Institutional Animal Care and Use Committee at Seattle Children’s Research Institute.

### Mouse immunization

Bacterial phage Qβ-derived virus-like particles (Qβ-VLPs) were generated as previously described (Liao et al., 2017) and provided by B. Hou (Chinese Academy of Sciences, Beijing, China). Mice were injected intraperitoneally with 2 μg Qβ-VLP-ssRNA. Analysis was performed 14 days after immunization.

### Flow cytometry

Tissues (spleen, lymph nodes, thymus, and lungs) were harvested into ice-cold FACS buffer (PBS with 2% FBS). Cells were resuspended in RBC lysis buffer (ACK), incubated for 2–3 min, and then spun down and resuspended in FACS buffer. Samples were then filtered through 70-μm cell strainers (Corning). Single-cell suspensions were stained with fluorescence-labeled antibodies for 20–30 min at 4°C. Intracellular staining was performed using the Foxp3 Staining Buffer Set (eBioscience) following cell-surface staining according to the manufacturer’s instructions. In some instances, cells were fixed with 2% PFA for 10 minutes for Foxp3 intracellular staining to preserve YFP. Lung lymphocytes were isolated from PBC perfused lung followed by followed by enzymatic digestion in the presence of 1.5-mg/ml collagenase type IV and 0.25 mg/ml DNaseI at 37°C in an incubator shaker for 45 minutes. The digested tissues were then filtered through a 40-μm filter and centrifuged. The cell pellet was resuspended in 5 ml of the 70% fraction of a 70:37 Percoll gradient and overlaid on 5 ml of the 37% fraction in a 15 ml Falcon tube (Genesee Scientific). Percoll gradient separation was performed by centrifugation without brakes for 20 minutes at 1500 rpm. Mononuclear cells were removed from the interphase, washed with cold PBS, and resuspended in culture medium for analysis.

The initial gating strategy performed included forward scatter (FSC)-A/side scatter (SSC)-A, exclusion of doublets (through SSC-H/SSC-W and FSC-H/FSC-W), and live cells. Cell counts were determined using Spherotech Accucount Fluorescent Beads (Thermo Fisher Scientific). After total cell calculation, the number of specific populations was determined by the frequency of the population determined by flow cytometry. Samples were acquired on a LSRII (BDBiosciences) flow cytometer, and data were analyzed using FlowJo software (Tree Star).

### Reagents

Anti-murine antibodies used in the study include the following: B220 (RA3-6B2), CD19 (1D3), CD138 (281–2), GL7 (GL7), CD38 (90), IgM (II/41), IgD (11–26), CD45 (30-F11), CD4 (GK1.5), CD8 (53-6.7), CD44 (IM7), CD62L (MEL-14), CD25 (PC61), CTLA-4 (UC10-4B9), and Helios (22F6) from Bio-Legend; IgG2c (polyclonal) from Southern Biotech; CD25 (PC61.5), PD-1 (J43), and IFN-γ (XMG1.2) from eBioscience/Thermo Fisher Scientific; ICOS (7E.17G9) and CXCR5 (2G8) from BD Biosciences. Qβ-specific cell labeling methods, as previously described (Liao et al., 2017) were used to to characterize the Ag-specific B cells.

### ddPCR assay

Treg (CD4^+^CD25^+^) were isolated from the spleen and lymph nodes of FoxP3-aPIK3CD mice using the CD4^+^CD25^+^ Regulatory T Cell Isolation Kit (Miltenyi Biotec). Genomic DNA was isolated from these cells using the AllPrep DNA/RNA Micro Kit (Qiagen). Digital droplet (dd)PCR assays were performed as previously described (Wray-Dutra et al., 2018).

### Measurement of autoantibodies

Autoantigen microarrays were performed at the University of Texas Southwestern Microarray Core as previously described (Li et al., 2007).

### ELISA

96-well Nunc-Immuno MaxiSorp plates (Thermo Fisher Scientific) were coated overnight at 4°C with 2 μg/ml VLP in PBS. Plates were blocked with 2% BSA before sample incubation, and then plates were incubated with serially diluted serum at RT for 2 h. Subsequently, plates were incubated with HRP conjugated goat anti-mouse IgM, IgG, or IgG2c (1:2,000 dilution; Southern Biotech). Peroxidase reactions were developed using OptEIA TMB substrate reagent set (BD Biosciences) and stopped with sulfuric acid. Absorbance was measured at 450 nm using a SpectraMax i3X microplate reader (Molecular Devices). Autoantigen microarrays were performed at the UT Southwestern Medical Center Microarray Core Facility, Dallas, TX (Li et al., 2007).

### Suppression assay

Suppression assays were performed by coculturing 1 × 10^5^ Cell Trace Violet (CTV) labelled CD4^+^CD25^-^ effector T cells (Teff) with varying ratios of CD4^+^CD25^+^ Treg isolated from either FoxP3-aPIK3CD or FoxP3-WT mice in the presence of 5 × 10^5^ irradiated APCs (autologous CD4^-^ splenocytes irradiated at 2500 rad) and 0.5ug/ml soluble anti-CD3. Dilution of CTV was measured by flow cytometry after 4 days of coculture. Dilution of CTV was interpreted as proliferation of Teff, and % suppression was calculated as (a − b)/a × 100, where a is the percentage of Teff proliferation in the absence of Treg, and b is the percentage of Teff proliferation in the presence of Treg.

### Statistical analysis

Statistical significance was determined with Prism9 (GraphPad) software Version 9.0/10 using two-tailed unpaired Student’s t-test or ANOVA multiple comparison as indicated in relevant figure legends.

## Acknowledgments

The authors thank J. Smith, S. Khim and A. Zielinska-Kwiatkowska for the technical assistance and mouse colony maintenance. This work was supported by National Institutes of Health awards DP3-DK111802 (D.J. Rawlings). Additional support was provided by the Children’s Guild Association Endowed Chair in Pediatric Immunology (D.J. Rawlings), the Tom Hansen Investigator in Pediatric Innovation Endowment (D.J. Rawlings), the Benaroya Family Gift Fund (D.J. Rawlings), and the Seattle Children’s Research Institute, Program for Cell and Gene Therapy.

## Author contributions

A.S., F.A.Q., and T.D. designed and performed experiments, analyzed data, and generated figures. B.H. provided VLP reagents and intellectual input into experimental design and interpretation of immunization studies. D.J.R. designed and supervised the study and interpreted data. A.S., F.A.Q., T.D., and D.J.R. wrote the manuscript. All the authors edited the manuscript.

---

**Figure 2–figure supplement 1. Spontaneous GC responses and autoantibodies in young Foxp3-aPIK3CD mice.** (A) Frequency and absolute number of GC B cells in the spleen of 12-wk-old Foxp3-WT and Foxp3-aPIK3CD mice. (B) Frequency and absolute number of CD138^+^ PCs in the spleen of 12-wk-old Foxp3-WT and Foxp3-aPIK3CD mice. (C) Autoantibody microarray heatmap shows signal intensity of IgM autoantibodies to the most significant autoantigens in the serum of young (12wks) and old (34wks) Foxp3-WT and Foxp3-aPIK3CD mice.

**Figure 5–figure supplement 2. Treg expressing aPIK3CD exhibit phenotypic changes including increased PD-1 levels.** A) Graphs showing the MFI of Treg associated proteins including Foxp3, CD25, CTLA-4, PD-1, and Helios in Treg isolated from thymus, spleen and lung of Foxp3-WT (black circles) and Foxp3-aPIK3CD mice (red circles). Two independent experiments were performed (n ≥ 3 mice per group). Significance determined by t-test (unpaired). *, P < 0.05 **, P < 0.01, Graphs depict mean.

**Figure 7–figure supplement 3. VLP-specific antibody titers in immunized Foxp3-aPIK3CD mice.** Dilution curves of VLP-specific IgM, IgG, and IgG2c in serum of 12- to 14-wk-old Foxp3-WT and Foxp3-aPIK3CD mice at day 14 d after intraperitoneal immunization with VLP-ssRNA.

