## Supplementary figures and images for "Activated PI3Kδ specifically perturbs mouse Treg homeostasis and function leading to immune dysregulation"

### Figure 2-figure supplement 1

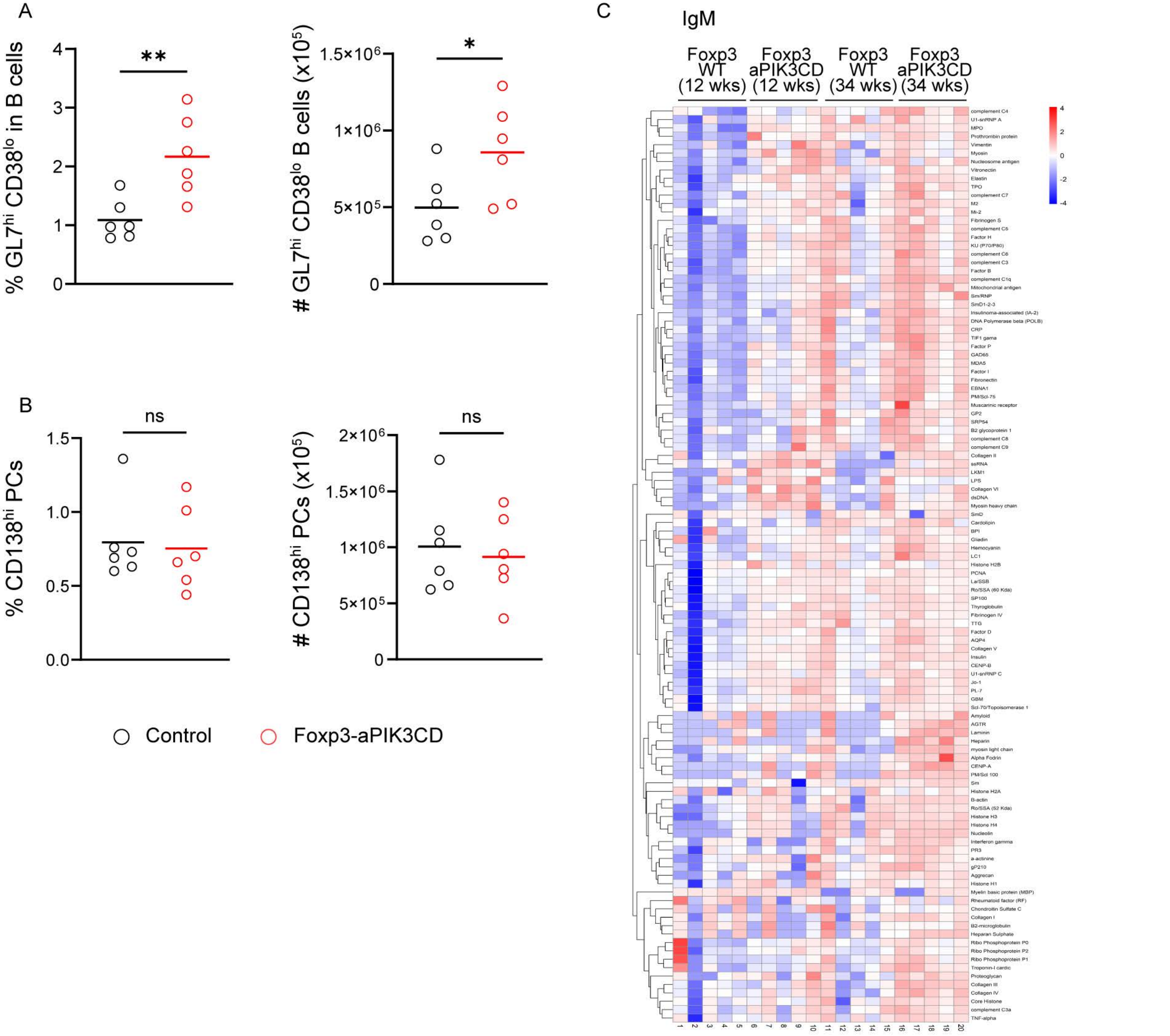

### Figure 5-figure supplement 2

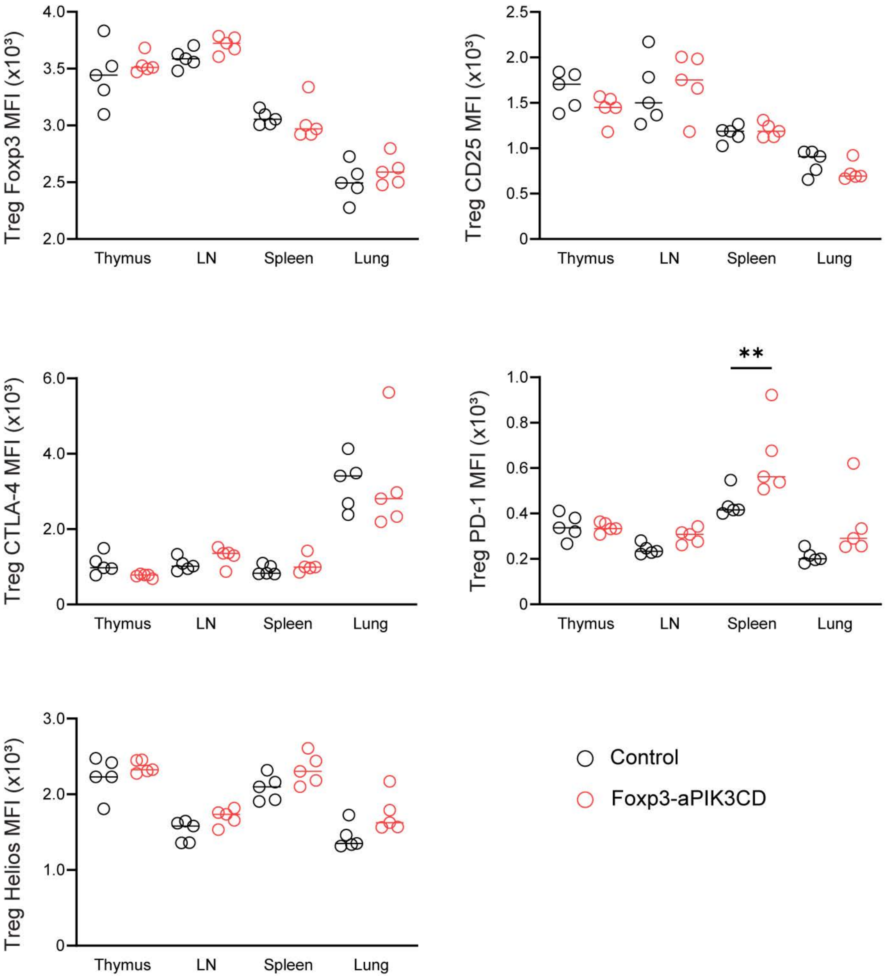

### Figure 7-figure supplement 3

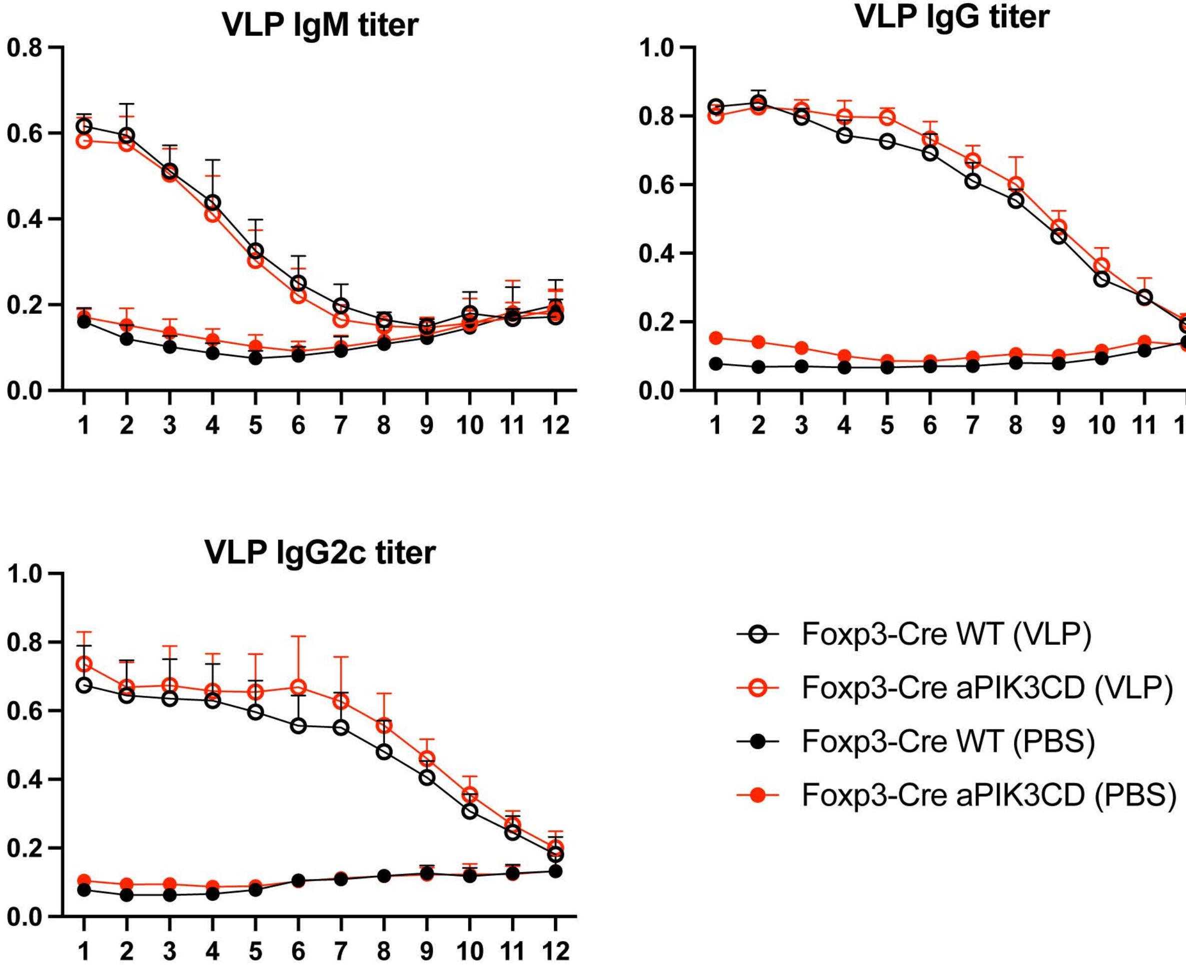
